# A thermostable Cas9 with increased lifetime in human plasma

**DOI:** 10.1101/138867

**Authors:** Lucas B. Harrington, David Paez-Espino, Janice S. Chen, Enbo Ma, Brett T. Staahl, Nikos C. Kyrpides, Jennifer Doudna

## Abstract

CRISPR-Cas9 is a powerful technology that has enabled genome editing in a wide range of species. However, the currently developed Cas9 homologs all originate from mesophilic bacteria, making them susceptible to proteolytic degradation and unsuitable for applications requiring function at elevated temperatures. Here, we show that the Cas9 protein from the thermophilic bacterium *Geobacillus stearothermophilus* (GeoCas9) catalyzes RNA-guided DNA cleavage over a wide temperature range and has an enhanced protein lifetime in human plasma. GeoCas9 is active at temperatures up to 70°C, compared to 45°C for *Streptococcus pyogenes* Cas9 (SpyCas9), which greatly expands the temperature range for CRISPR-Cas9 applications. By comparing features of two closely related *Geobacillus* homologs, we created a variant of GeoCas9 that doubles the DNA target sequences that can be recognized by this system. We also found that GeoCas9 is an effective tool for editing mammalian genomes when delivered as a ribonucleoprotein (RNP) complex. Together with an increased lifetime in human plasma, the thermostable GeoCas9 provides the foundation for improved RNP delivery *in vivo* and expands the temperature range of CRISPR-Cas9.

---

The use of CRISPR-Cas9 has rapidly transformed the ability to edit and modulate the genomes of a wide range of organisms^1^. This technology, derived from adaptive immune systems found in thousands of bacterial species, relies on RNA-guided recognition and cleavage of invasive viral and plasmid DNA^2^. The Cas9 proteins from these species differ widely in their size and cleavage activities^3–5^. Despite the abundance and diversity of these systems, the vast majority of applications have employed the first Cas9 homolog developed from *Streptococcus pyogenes* (SpyCas9)^6^. In addition to SpyCas9, several other Cas9 and related proteins have also been shown to edit mammalian genomes with varying efficiencies^5,7–10^. While these proteins together provide a robust set of tools, they all originate from mesophilic hosts, making them unsuitable for applications requiring activity at higher temperatures or extended protein stability.

This temperature restriction is particularly limiting for genome editing in obligate thermophiles^11^. Recent efforts using SpyCas9 to edit a facultative thermophile have been possible by reducing the temperature within the organism^12^. While effective, this approach is not feasible for obligate thermophiles, and requires additional steps for moderate thermophiles. This is especially important for metabolic engineering for which thermophilic bacteria present enticing hosts for chemical synthesis due to decreased risk of contamination, continuous recovery of volatile products and the ability to conduct reactions that are thermodynamically unfavorable in mesophilic hosts^13^. Developing a thermostable Cas9 system will enable facile genome editing in thermophilic organisms using technology that is currently restricted to mesophiles.

CRISPR-Cas9 has also emerged as a potential treatment for genetic diseases^14^. A promising method for the delivery of Cas9 into patients or organisms is the injection of preassembled Cas9 ribonucleoprotein complexes (RNP) into the target tissue or bloodstream^15^. One major challenge to this approach is that Cas9 must be stable enough to survive degradation by proteases and RNases in the blood or target tissue for efficient delivery. Limited protein lifetime will require delivery of higher doses of Cas9 into the patient or result in poor editing efficacy. In contrast, delivering a Cas9 with improved stability could greatly enhance genome-editing efficiency *in vivo*.

To address these challenges, we tested the thermostable Cas9 protein from *Geobacillus stearothermophilus* (GeoCas9). We find that GeoCas9 maintains activity over a wide temperature range. By harnessing the natural sequence variation of GeoCas9 from closely related species, we engineered a protospacer adjacent motif (PAM)-binding variant that recognizes additional PAM sequences and thereby doubles the number of targets accessible to this system. We also engineered a highly efficient single-guide RNA (sgRNA) based on RNAseq data from the native organism and show that GeoCas9 can efficiently edit genomic DNA in mammalian cells. The functional temperature range of GeoCas9 complements that of previously developed Cas9 systems, greatly expanding the utility and stability of Cas9 for both *in vitro* cleavage and genome editing applications.

## Results

### Identification of thermostable Cas9 homologs

Although thousands of *cas9* homologs have been sequenced, there have been no functionally validated Cas9 orthologs from archaea^16^, restricting our search for a thermophilic Cas9 to thermophilic bacteria. We examined all the isolates in Integral Microbial Genomes database (IMG) from a thermophilic environment that contained a Cas9-like protein^17^ (hits to a TIGRfam model 01865 for Csn1-like or 03031 for a Csx12-like). From this analysis, the Cas9 from *Geobacillus stearothermophilus* (*G. st.*; formerly *Bacillus stearothermophilus*)^18^ stood out because it was full-length and its sequence is shorter than that of the average Cas9. Most importantly, this candidate is found in an = organism that can grow at a reported temperature range of 30°C–75°C (optimal at 55°C). A BLASTn search for proteins similar to GeoCas9 revealed several nearly identical homologs (from 93.19–99.91% identity over the full length) in six other *Geobacillus* species (ED Table 2) and 92.55% Identity over the full length in *Effusibacillus pohliae* DSM22757. *G. st.* has been a proven source of enzymes for thermophilic molecular cloning applications^19^, thermostable proteases^20^ and enzymes for metabolic engineering^21^. Moreover, the wide temperature range that *G. st.* occupies^22^ suggested that the Cas9 from this species (GeoCas9) might able to maintain activity at both mesophilic and thermophilic temperatures (Fig. 1a). Notably, GeoCas9 is considerably smaller than SpyCas9 (GeoCas9, 1,087 amino acids; SpyCas9, 1,368 amino acids). A homology model of GeoCas9 based on available Cas9 crystal structures along with sequence alignments revealed that the small size of GeoCas9 is largely the result of a reduced REC lobe, as is the case with other compact Cas9 homologs from *Streptococcus aureus* Cas9 (SauCas9) and *Actinomyces naeslundii* Cas9 (AnaCas9) (Fig. 1b, c; Supplementary Fig. 1).

**Figure 1.**
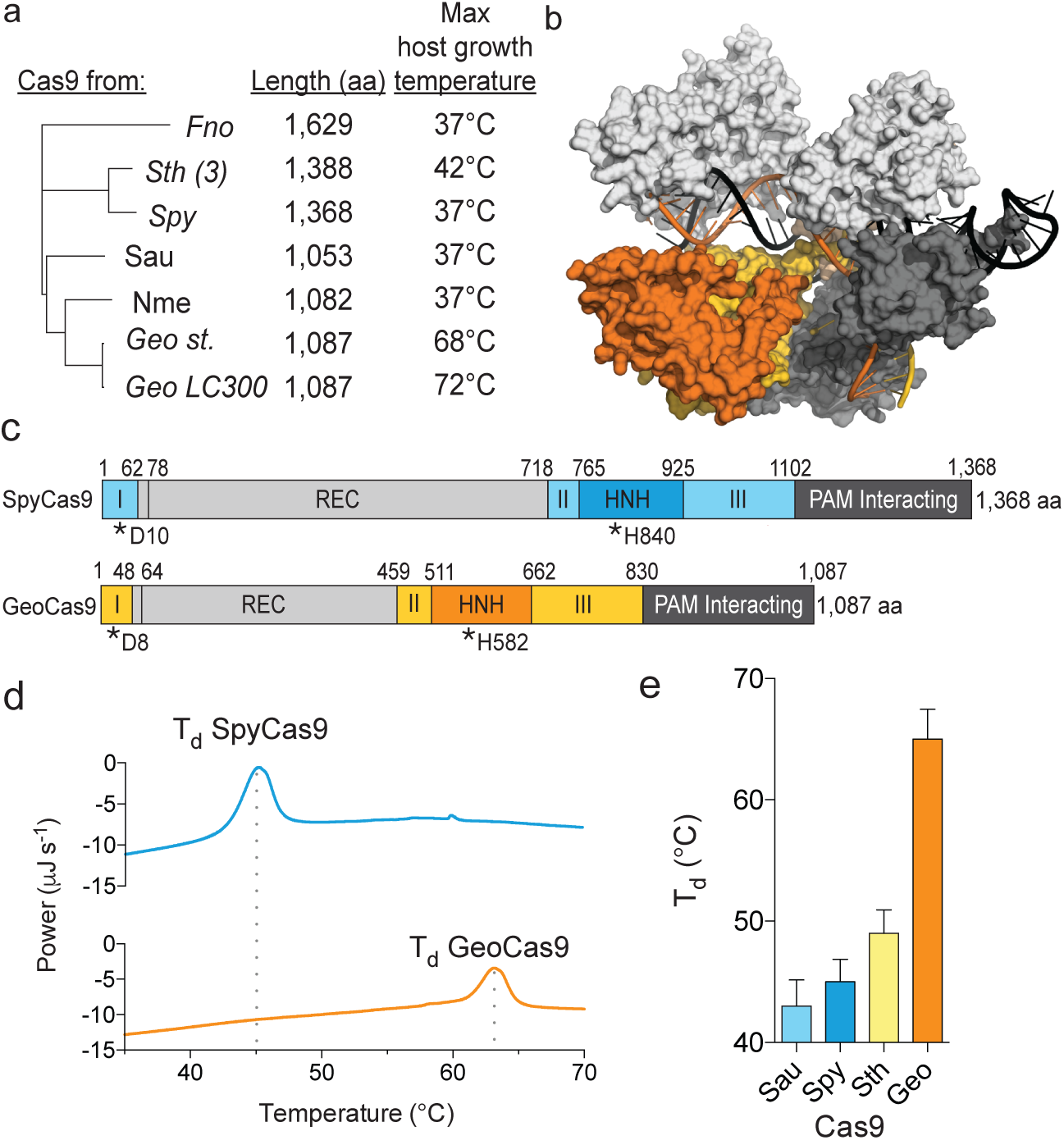
GeoCas9 is a small, thermostable Cas9 homolog. **a**, Phylogeny of Cas9 proteins used for genome editing with their length (amino acids) and the maximum temperatures that supports growth of the host indicated to the right ^22^ (Nme*, Neisseria meningitidis*; Geo, *Geobacillus stearothermophilus*; Geo LC300, *Geobacillus* LC300; Spy*, Streptococcus pyogenes;* Sau, *Streptococcus aureus*; Fno, *Francisella novicida*; Sth (3), *Streptococcus thermophilus* CRISPR III). **b**, Homology model of GeoCas9 generated using Phyre 2^42^ with the DNA from PDB 5CZZ docked in. **c**, Schematic illustration of the domains of Spy Cas9 (blue) and GeoCas9 (orange) with active site residues indicated below with asterisks. **d**, Representative traces for Differential Scanning Calorimetry (DSC) of GeoCas9 and SpyCas9, T_d_; Denaturation temperature. **e,** Denaturation temperature of various Cas9 proteins as measured by DSC, mean ± S.D. is shown.

We purified GeoCas9 and performed initial thermostability tests using differential scanning calorimetry (DSC), which showed that in the absence of RNA or DNA, GeoCas9 has a denaturation temperature about 20°C higher than SpyCas9 (Fig. 1d). Moreover, GeoCas9 denatures at 15°C higher than the slightly thermophilic *Streptococcus thermophilus* CRISPR III Cas9 (SthCas9) (Fig. 1e). Given these results, we selected GeoCas9 as a candidate for further development and optimization.

### GeoCas9 PAM identification and engineering

CRISPR systems have evolved a preference for a protospacer adjacent motif (PAM) to avoid self-targeting of the host genome^23,24^. These PAM sequences are divergent among Cas9 homologs and bacteriophage DNA targets are often mutated in this region to escape cleavage by Cas9^25^. To identify the PAM for GeoCas9, we first searched for naturally targeted viral and plasmid sequences using CRISPRtarget^26^. The three sequenced strains of *G. st*. provided 77 spacer sequences, and 3 of them had high-confidence viral and plasmid targets (Supplementary Fig. 2, Supplementary Table 3,). Extracting the sequences 3′ of the targeted sequence revealed a consensus of 5′-NNNNCNAA-3′ (Fig. 2a, ED Fig. 2). Given the low number of viral targets, we next performed cleavage assays on substrates containing various PAM sequences, revealing a complete PAM sequence of 5′-NNNNCRAA-3′ (Fig. 2b).

**Figure 2.**
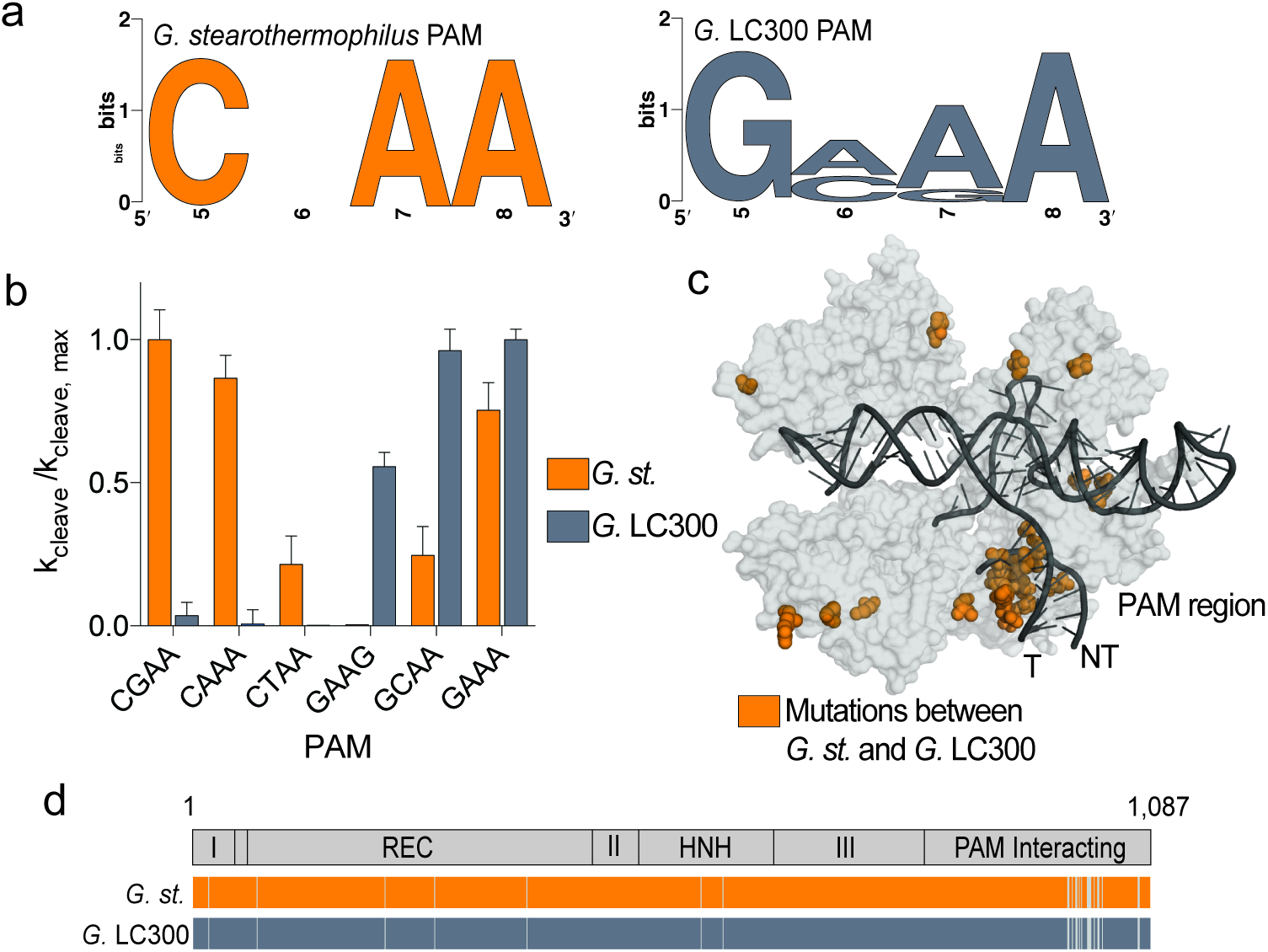
PAM identification and engineering of GeoCas9. **a**, WebLogo for sequences found at the 3′ end of protospacer targets identified with CRISPRTarget for *Geobacillus stearothermophilus* (left panel) and *Geobacillus* LC300 (right panel). **b**, Cleavage assays conducted with the two homologs of GeoCas9. Substrates with various PAM sequences were P32-labelled and mean ± S.D. is shown. **c**, Mapping of mutated residues (orange spheres) between *G. st.* and *G.* LC300 onto the homology model of GeoCas9 showing high density in the PAM interacting domain near the PAM region of the target DNA. **d**, Alignment of the Cas9 proteins from *G. st.* and *G.* LC300 with the domain boundaries shown above. Solid colors represent identical residues and grey lines indicate residues that are mutated between the two Cas9 homologs.

In addition to the CRISPR loci found in *G. st.* strains, we also found a type II CRISPR locus in *Geobacillus* LC300 containing a Cas9 with ~97% amino acid identity to the *G. st.* Cas9. Despite having nearly identical sequences, alignment of these two homologs of GeoCas9 revealed a tight cluster of mutations in the PAM interacting domain (PI). Furthermore, mapping these mutations onto the homology model of GeoCas9 showed that they are located near the PAM region of the target DNA (Fig. 2c, d). We hypothesized that this GeoCas9 variant might have evolved altered PAM specificity. By searching for viral targets using the spacers in the *G.* LC300 array, we identified a preference for GMAA in place of the CRAA PAM of *G. st.*, lending support to our hypothesis. We constructed and purified a hybrid Cas9 protein in which the PI domain of the *G.* LC300 Cas9 was substituted for the PI domain of *G. st.* Cas9 and tested cleavage activity on targets containing various PAM sequences (Fig. 2b). We found that, as predicted by protospacer sequences, the hybrid Cas9 preferred a GMAA PAM rather than the CRAA PAM utilized by GeoCas9. Moreover, *G.* LC300 appears to be more specific for its optimal PAM, which may result in lower off-target cleavage for genome editing applications^27^. By creating a hybrid Cas9 with this naturally occurring PAM-recognition variant, we double the sequence space that can be targeted by GeoCas9 without resorting to structure based protein engineering as has been done for other Cas9 homologs^27^.

### Identification of both crRNA and tracrRNA and engineering of GeoCas9 sgRNA

CRISPR-Cas9 systems use a trans-activating crRNA (tracrRNA), which is required for maturation of the crRNA and activation of Cas9^6,28^. To identify the tracrRNA for GeoCas9, we cultured *G. st.* and deep sequenced the small RNA it produced. We found that the CRISPR array was transcribed despite a lack of phage or plasmid challenge, and that the array was transcribed in the opposite direction of the Cas proteins (Fig. 3a). The crRNA was processed to 23nt (Fig. 3b) of the spacer sequence and 18nt of the repeat sequence *in vivo*, similar to other small type IIC Cas9 proteins^8,29^. Mapping of the RNAseq reads to the CRISPR array also revealed a putative tracrRNA upstream of the Cas9 open reading frame (ORF).

**Figure 3.**
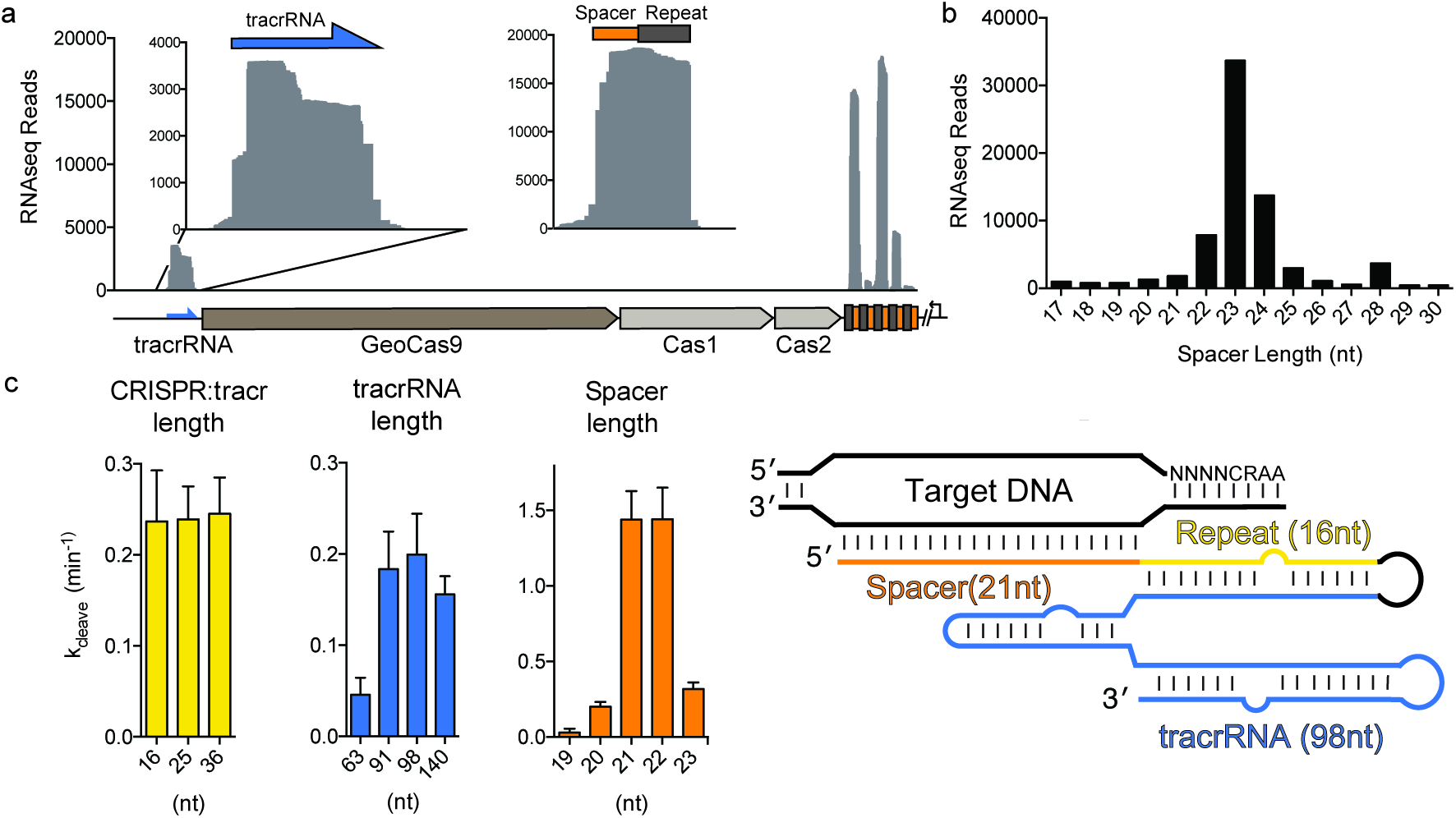
Small RNA-seq and sgRNA engineering for GeoCas9. **a**, Small RNA sequenced from *G. stearothermophilus* mapped to the CRISPR locus. Inset shows enlargement of the region corresponding to the tracrRNA and the most highly transcribed repeat and spacer sequence. **b**, Distribution of the length of the spacer sequences extracted from the small RNA sequencing results. **c**, Length optimization of the tracrRNA and crRNA for GeoCas9 and the optimal guide RNA design (right panel). The length of the tracrRNA, crRNA:tracrRNA duplex and spacer was optimized sequentially by transcribing variations of the sgRNA and testing their ability to guide GeoCas9-mediated cleavage of a radiolabeled substrate. The mean k_cleave_ ± S.D. is shown and experiments were conducted in triplicate.

We joined this putative tracrRNA to the processed crRNA using a GAAA-tetraloop to generate a single-guide RNA (sgRNA)^30^. Variations of this sgRNA were *in vitro* transcribed and tested for their ability to direct GeoCas9 to cleave a radiolabeled double-stranded DNA target at 37°C. We first varied the length of the crRNA:tracrRNA duplex and found that this modification had little impact on the DNA cleavage rate (left panel, Fig. 3c), making it a valuable place for further sgRNA modifications^31^. Next, we tested the length of the tracrRNA, choosing stopping points near predicted rho-independent terminators. In contrast to the crRNA:tracrRNA duplex length, the length of the tracrRNA had a dramatic effect on the cleavage rate, with sequences shorter than 91nt supporting only a small amount of cleavage (middle panel, Fig. 3c). Finally, we varied the length of the spacer sequence and found that 21–22nt resulted in a more than 5-fold increase in cleavage rate, compared to the 20nt spacer preferred by SpyCas9 (right panel, Fig. 3c). This finding contrasts with the most abundant spacer length of 23nt found by RNAseq. This difference may be due to inter- or intramolecular guide interactions in the *in vitro* transcribed sgRNA^32^.

### Genome editing by GeoCas9 RNPs in mammalian cells

With evidence that GeoCas9 maintains cleavage activity at mesophilic temperatures, we assessed the ability of GeoCas9 to edit mammalian genomes. We compared GeoCas9 and SpyCas9 editing efficiency by delivering preassembled ribonucleoprotein complexes (RNPs) into cultured cells, circumventing differences between SpyCas9 and GeoCas9 protein expression. First, GeoCas9 RNPs targeting regions adjacent to various PAM sequences were delivered into HEK293T cells expressing a destabilized GFP (Fig. 4a). We found that when targeted to sequences adjacent to the preferred CRAA PAM, GeoCas9 decreased GFP fluorescence at levels comparable to those observed for SpyCas9 (Fig. 4a). Next, we targeted GeoCas9 to cleave the native genomic loci DNMT1 and AAVS1 (Fig. 4b,c). We varied the length of the targeting spacer sequence and found that at one site 21nt was a sufficient length to efficiently induce indels while at another site a 22nt spacer length was necessary. Given this variability and that extending the spacer length to 22nt had no detrimental effects, we conclude that a 22nt guide segment length is preferred for use in genome editing applications. Moreover, when we tested editing efficiency at a site containing an overlapping PAM for both GeoCas9 and SpyCas9, we observed similar editing efficiencies by both proteins (Fig. 4b). At the DNMT1 locus we titrated amounts of GeoCas9 and SpyCas9 RNPs to assess the effect on genome editing efficiency (Fig. 4c). Products analyzed by T7E1 assay again showed efficient production of indels by both GeoCas9 and SpyCas9. These results demonstrate that GeoCas9 is an effective alternative to SpyCas9 for genome editing in mammalian cells.

**Figure 4.**
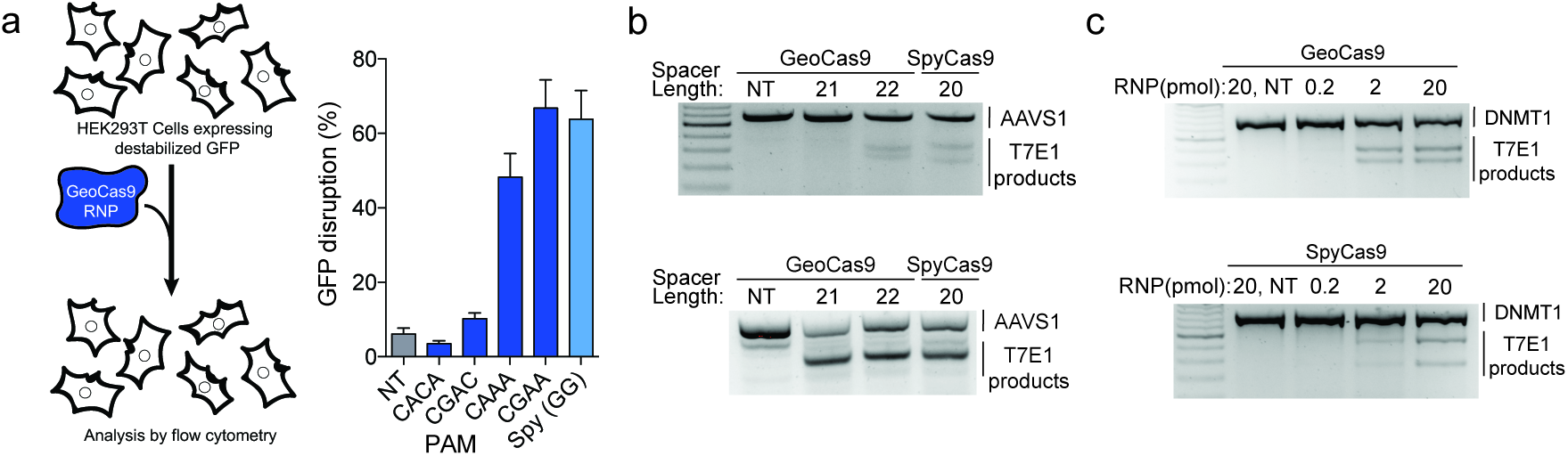
Genome editing activity of GeoCas9 in mammalian cells. **a**, EGFP disruption in HEK293T cells by GeoCas9. HEK293FT cells expressing a destabilized GFP were transfected with GeoCas9 RNP preassembled with a targeting or non-targeting guide RNA. Cells were analyzed by flow cytometry and targets adjacent to the CRAA PAM resulted in efficient GFP disruption (NT; non-targeting). **b**, T7E1 analysis of indels produced at the AAVS1 locus when the guide length was varied from 21 to 22nt. **c**, T7E1 analysis of indels produced using a titration of GeoCas9 and SpyCas9 RNP targeting the DNMT1 locus in HEK293T cells.

### GeoCas9 functions over a wide temperature range and has extended lifetime in human plasma

Based on initial observations showing that GeoCas9 protein remains folded at elevated temperatures (Fig. 1d, e), we tested whether the GeoCas9 RNP maintains activity after exposure to high temperatures. We incubated SpyCas9 and GeoCas9 at a challenge temperature and added equimolar substrate to test the fraction of RNP that remained functional. After incubation for 10 min at 45°C, the fraction of active SpyCas9 was greatly reduced (Fig. 5a). In contrast, the fraction of GeoCas9 after incubation at 45°C remained at 100% and not until challenge at 70°C did we detect a decrease in activity (Fig. 5a).

**Figure 5.**
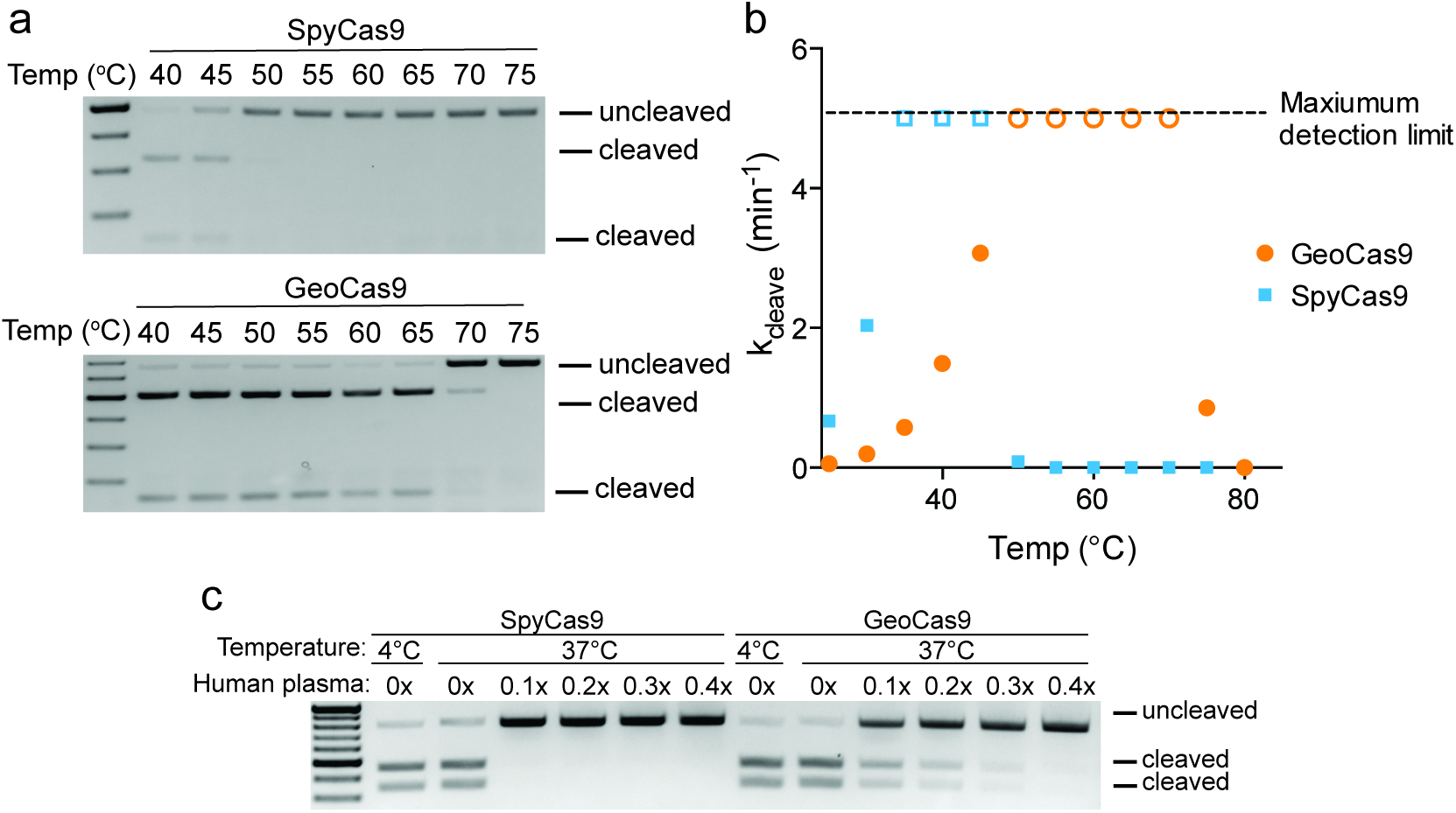
Thermostability of GeoCas9 and longevity in human plasma. **a**, Activity of SpyCas9 and GeoCas9 after incubation at the indicated temperature. After challenging at the higher temperature, reactions were conducted at 37°C using a 1:1 ratio of substrate to RNP. **b**, Cleavage rate of SpyCas9 and GeoCas9 RNPs at various temperatures. Maximum detection limit is shown by the dashed line at k_cleave_=5, indicating that the reaction completed in ≤30s. **c**, Effect of incubating GeoCas9 and SpyCas9 in human plasma. After incubation in varying concentrations of human plasma for 8hrs at 37°C, the reaction was carried out with 1:1 ratio of DNA substrate to RNP.

Often thermostability comes at the cost of reduced activity at lower temperatures^33^. However, the wide range of natural growth temperatures for *G. st.* suggested that GeoCas9 might maintain activity at mesophilic temperatures. To examine this hypothesis, we measured the cleavage rate of SpyCas9 and GeoCas9 at various temperatures (Fig. 5b). SpyCas9 DNA cleavage rates increased between 20–35°C, reaching maximum levels from 35–45°C. Above these temperatures, SpyCas9 activity dropped sharply to undetectable levels, as predicted by thermostability measurements. In contrast, GeoCas9 activity increased to a maximum measured value at 50°C and maintained maximum detectable activity up to 70°C, dropping to low levels at 75°C. These results make GeoCas9 a valuable candidate for editing obligate thermophilic organisms and for biochemical cleavage applications requiring Cas9 to operate at elevated temperatures.

It was shown previously that thermostable proteins have longer lifetime in blood^34^. To test if this is the case for GeoCas9, we incubated SpyCas9 and GeoCas9 in diluted human plasma at 37°C for 8 hrs and measured the amount of Cas9 activity remaining (Fig. 5c). Although SpyCas9 maintained activity when incubated in reaction buffer at 37°C, its activity was abolished even at the lowest concentration of plasma. In contrast, GeoCas9 maintained robust activity after incubation with human plasma, making it a promising candidate for *in vivo* RNP delivery.

## Discussion

Our results establish GeoCas9 as a thermostable Cas9 homolog and expand the temperatures at which Cas9 can be used. We anticipate that the development of GeoCas9 will enhance the utility of CRISPR-Cas9 technology at both mesophilic and thermophilic temperatures. The ability of Cas9 to function reliably in a wide range of species has been key to its rapid adoption as a technology, but the previously developed Cas9 homologs are limited for use in organisms that can grow below 42°C. The complementary temperature range of GeoCas9 with SpyCas9 (Fig. 5c) opens up Cas9– based genome editing to obligate thermophiles and facultative thermophiles, without the additional steps of altering the temperature of the organisms. We also anticipate that GeoCas9 will be useful for *in vitro* molecular biology applications requiring targeted cleavage at elevated temperatures. Furthermore, we predict that the extended lifetime of GeoCas9 in human plasma may enable more efficient delivery of Cas9 RNPs.

We were interested to note that GeoCas9 and SpyCas9 induced similar levels of indels in HEK293T cells as SpyCas9 when delivered as an RNPs (Fig. 4b-d), despite GeoCas9’s lower DNA cleavage rate at 37°C (Fig. 5b). We conclude that biochemical cleavage rates may not reflect limiting step of target search in a human cell. It may be that GeoCas9 can persist longer in cells, which raises its effective concentration over time and compensates for its slower cleavage rate. Moreover, in applications requiring delivery of Cas9 into the bloodstream, the benefit of improved stability by GeoCas9 may become even more apparent.

The development of GeoCas9 hinged upon utilizing the naturally occurring diversity of CRISPR systems. The sheer abundance and diversity of Cas9 makes it advantageous over newer type V systems, such as Cpf1, for developing specialized genome editing tools. It has previously been suggested that type II CRISPR systems are only found in mesophilic bacteria, and that protein engineering would be required to develop a thermophilic Cas9^35^. The rarity of type II CRISPR systems in thermophiles is surprising given that CRISPR systems in general are enriched in thermophilic bacteria and archaea^36^. However, by searching the continually growing number of sequenced bacteria, we uncovered a naturally occurring thermophilic Cas9. Exploiting Cas9 sequence diversity, rather than engineering thermostability, revealed a protein that maintains activity over a broad temperature range, which is often difficult to select for using directed evolution. Using the natural context of this CRISPR system, including the transcribed RNA and targeted sequences, we further developed GeoCas9 with minimal experimental optimization. The strategy of mining the natural context and diversity of CRISPR systems has proven successful for uncovering novel interference proteins^16,37^, and we anticipate that it can be applied more broadly to discover and develop new genome editing tools.

## Methods

### Identification of thermophilic Cas9 homologs and generation of heterologous expression plasmids

We mined all isolate genomes from the public Integrated Microbial genomes (IMG) database^17^ using the “Genome Search by Metadata Category tool.” We selected all the genomes annotated as “thermophile” (336) or “hyperthermophile” (94) and searched for the presence of Cas9-like candidates (hits to a TIGRfam model 01865 for Csn-like or 03031 for a Csx12-like) contained within a full CRISPR-Cas system (presence of Cas1, Cas2, and a Repeat-Spacers array). We initially selected the GeoCas9 variant due to its completeness, smaller gene size (shorter than the widely used SpyCas9), and growth in a large temperature range from 30-75 (optimal at 55C).The Cas9 from *Geobacillus stearothermophilus* was codon optimized for *E. coli*, ordered as Gblocks (IDT) and assembled using Gibson Assembly. For protein expression, a pET based plasmid containing an N terminal 10xHis-tag and MBP was used. For PAM depletion assays, a p15A plasmid was generated with the sgRNA constitutively expressed. The plasmids used are available from Addgene and their maps can be found in Supp. Table 2.

### Cas9 purification

Cas9 was purified as previously described^6^ with modification. After induction, *E. coli* BL21(DE3) expressing Cas9 was grown in Terrific Broth overnight at 18°C. Cells were harvested, re-suspended in Lysis Buffer (50mM Tris-HCl, pH 7.5, 20mM imidazole, 0.5mM TCEP, 500mM NaCl, 1mM PMSF), broken by sonication, and purified on Ni-NTA resin. TEV was added to the elution and allowed to cleave overnight at 4°C. The resulting protein was loaded over tandem columns of an MBP affinity column onto a heparin column and eluted with a linear gradient from 300mM to 1250mM NaCl. The resulting fractions containing Cas9 were purified by gel filtration chromatography and flash frozen in Storage Buffer (20mM HEPES-NaOH pH 7.5, 5% Glycerol, 150mM NaCl, 1mM TCEP).

### Differential Scanning Calorimetry

Cas9 proteins were dialyzed into degassed DSC Buffer (0.5mM TCEP, 50mM KH_2_PO_4_(pH 7.5), 150mM NaCl) overnight at 4°C. Samples were diluted to 0.3mg/ml and loaded a sample cell of a NanoDSC (TA instruments); buffer alone was used in the reference cell. The cell was pressurized to 3atm and the sample was heated from 20 to 90°C. Measurements made for buffer in both the sample and reference cells were subtracted from the sample measurements.

### Biochemical cleavage assays

Radioactive cleavage assays were conducted as previously described^38^. Reactions were carried out in 1× Reaction Buffer (20mM Tris-HCl, pH 7.5, 100mM KCl, 5mM MgCl_2_, 1mM DTT and 5% glycerol (v/v)). 100nM Cas9 and 125nM sgRNA were allowed to complex for 5min at 37°C. ~1nM radiolabeled probe was added to the RNP to initiate the reaction. Finally, the reaction was quenched with 2× Loading Buffer (90% formamide, 20mM EDTA, 0.02% bromophenol blue, 0.02% xylene cyanol and products were analyzed on 10% urea-PAGE gel containing 7M urea.

For thermostability measurements (Fig. 4a), 100nM Cas9 was complexed with 150nM sgRNA in 1x Reaction Buffer for 5min at 37°C. 100nM of a PCR product containing the targeted sequence was cleaved using dilutions of the estimated 100nM RNP complex to accurately determine a 1:1 ratio of Cas9 to target. Next, samples were challenged at the indicated temperature (40°C–75°C) for 10min and then returned to 37°C. 100nM PCR product containing the targeted sequence was added to the reaction and it was allowed to react for 30min at 37°C. The reaction was quenched with 6× Quench Buffer (15% glycerol (v/v), 1mg/ml Orange G, 100mM EDTA) and products were analyzed on a 1.25% agarose gel stained with ethidium bromide.

For thermophilicity measurements (Fig. 4b), 500nM Cas9 was complexed with 750nM sgRNA in 1× Reaction Buffer for 5min at 37°C. The samples were placed at the assayed temperature (20°C–80°C) and 100nM of PCR product was added to the reaction. Time points were quenched using 6× Quench Buffer and analyzed on a 1.25% agarose gel stained with SYBR Safe (Thermo Fisher Scientific).

To study the effect of human plasma on the stability of Cas9 proteins, preassembled Cas9-RNP was incubated for 8 hours either at 37°C or 4°C in Reaction Buffer with the specified amount of normal human plasma. Substrate was then added and cleavage products were analyzed as described for thermostability measurements.

### Small RNA sequencing

*Geobacillus stearothermophilus* was obtained from ATCC and cultured at 55°C in Nutrient Broth (3g beef extract and 5g peptone per liter water, pH 6.8) to saturation. Cells were pelleted and RNA was extracted using a hot phenol extraction as previously described^39^. Total RNA was treated with TURBO DNase (Thermo Fisher Scientific), rSAP (NEB) and T4 PNK (NEB) according to manufactures instructions. Adapters were ligated onto the 3’ and 5’ ends of the small RNA, followed by reverse transcription with Superscript III. The library was amplified with limited cycles of PCR, gel-extracted on an 8% native PAGE gel and sequenced on an Illumina MiSeq. Adapters were trimmed using Cutadapt and sequences >10nt were mapped to the G. st. CRISPR locus using Bowtie 2^40^.

### HEK293T EGFP disruption assay and indel analysis

HEK293T cells expressing a destabilized GFP were grown in Dulbecco’s Modification of Eagle’s Medium (DMEM) with 4.5g/L glucose L-glutamine and sodium pyruvate (Corning Cellgro), supplemented with 10% fetal bovine serum, penicillin and streptomycin at 37°C with 5% CO_2_. ~24hrs before transfection, ~3 × 10^4^ cells were seeded into each well of a 96-well plate. The next day, 20pmol (unless otherwise specified) of RNP was assembled as previously described^41^ and mixed with 10µL OMEM. The RNP was added to 10µl of 1:10 dilution of Lipofectamine 2000 (Life Technologies) in OMEM and allowed to incubate at room temperature for 10min and added to the cells. Cells were analyzed for GFP fluorescence 48h later using Guava EasyCyte 6HT. Experiments were conducted in triplicate and the mean ± S.D. is shown. For analysis of indels, genomic DNA was extracted using Quick Extraction Solution (Epicentre), and the DNMT1 and AAVS1 loci were amplified by PCR. T7E1 reaction was conducted according to the manufacturer’s instructions and products were analyzed on a 1.5% agarose gel stained with SYBR gold (Thermo Fisher Scientific).

---

Supplementary Table 1| DNA and RNA sequences used in this study

Supplementary Table 2| Thermophilic Cas9 candidates

Supplementary Table 3| Spacers identified from *Geobacillus stearothermophilus* and *Geobacillus* LC300

## Acknowledgments

We thank A. Tambe, N. Ma and K. Zhou for technical assistance and discussions. L.B.H and J.S.C are supported by a US National Science Foundation Graduate Research Fellowship. J.A.D. is an Investigator of the Howard Hughes Medical Institute. This research was supported in part by the Allen Distinguished Investigator Program, through The Paul G. Allen Frontiers Group and the National Science Foundation (MCB-1244557 to J.A.D.).

## Author Contributions

L.B.H. designed and executed experiments with help from J.S.C., E.M., B.S and J.A.D.; The search for thermophilic Cas9 homologs was conducted by D.P.E. and N. C. K. All authors revised and agreed to the manuscript.

## Materials & Correspondence

Plasmids used in this study are available on Addgene (#87700, #87703). RNA sequencing data will be available on the NCBI Sequence Read Archive (SRA). Correspondence and requests for materials should be addressed to J.A.D.

